# Independent evolution of polymerization in the Actin ATPase clan regulates hexokinase activity

**DOI:** 10.1101/686915

**Authors:** Patrick R Stoddard, Eric M. Lynch, Daniel P. Farrell, Quincey A. Justman, Annie M. Dosey, Frank DiMaio, Tom A. Williams, Justin M. Kollman, Andrew W. Murray, Ethan C. Garner

**Author notes:** co-corresponding authors. Correspondence should be addressed to (AWM) and (ECG).

## Abstract

The actin protein fold is found in cytoskeletal polymers, chaperones, and various metabolic enzymes. Many actin-fold proteins, like the carbohydrate kinases, do not polymerize. We find that Glk1, a *Saccharomyces cerevisiae* glucokinase, forms two-stranded filaments with unique ultrastructure, distinct from that of cytoskeletal polymers. In cells, Glk1 polymerizes upon sugar addition and depolymerizes upon sugar withdrawal. Glk1 polymerization inhibits its enzymatic activity, thus the Glk1 monomer-polymer equilibrium sets a maximum rate of glucose phosphorylation regardless of Glk1 concentration. A mutation eliminating Glk1 polymerization alleviates concentration-dependent enzyme inhibition, causing glucokinase activity to become unconstrained. Polymerization-based regulation of Glk1 activity serves an important function *in vivo*: yeast containing non-polymerizing Glk1 are less fit when growing on sugars and more likely to die when refed glucose. Glucokinase polymerization arose within the ascomycete fungi and is conserved across a group of divergent (150-200 mya) yeast. We show that Glk1 polymerization arose independently from other actin-related filaments and allows yeast to rapidly modulate glucokinase activity as nutrient availability changes.

**One-sentence summary:** Yeast glucokinase activity is limited by its polymerization, which is critical for cell viability during glucose refeeding.

## Introduction

The Actin ATPase clan (*1*) is a diverse group of structurally similar protein families found in all domains of life (*2*). While several of the Actin ATPase families form polymers, polymerization has not been observed in those acting as metabolic enzymes, such as hexose kinases, or the Hsp70/DnaK family of chaperones (*3*). There are two possible explanations for the presence of several families of cytoskeletal polymers in the Actin ATPase clan: either polymerization evolved several times independently, or all of the polymer-forming families descend from a single, ancient, polymerizing ancestor. Phylogenetic (*4*) and structural studies (*5*) support a single origin of polymerization.

Cells use a variety of mechanisms to change enzyme activity rapidly in response to changes in internal and external conditions. These include allosteric and post-translational regulation as well as changes in the physical state of proteins, i.e. forming conglomerates by self-assembling into filaments or phase-separated gels. Self-assembly can serve as a sensitive and tunable way to control enzyme activity if polymerization depends on the concentration of enzymes, their substrates, and/or their products. Enzyme polymerization has been proposed to regulate flux through reaction pathways (*6*), to store enzymes during starvation (*7*), and to measure and signal cellular states (*8*).

Hexokinases and glucokinases of fungi are from a single family (the hexokinase family) within the Actin ATPase clan, and are both more similar to mammalian hexokinases than mammalian glucokinases. We find that Glk1, a glucokinase in the budding yeast *S. cerevisiae*, forms filaments when bound to its substrates or products. Glk1 filaments lack the conserved interaction geometries observed in all other actin-related filaments, rather they show novel inter-monomer contacts. Evolutionary and structural analysis indicates Glk1’s ability to polymerize evolved independently from that of actin or other actin-related filaments. Glk1 polymerization acts as an activity clamp, setting an upper limit to the catalytic rate of the entire Glk1 pool. This activity limit protects starved cells from the toxic effect of sudden exposure to high levels of glucose: eliminating Glk1’s ability to polymerize results in increased cell death during this transition.

## Results

### Substrate binding induces polymerization of Glk1

In *S. cerevisiae*, the first step in glycolysis, glucose phosphorylation, is catalyzed by three enzymes: two hexokinases (Hxk1 and Hxk2) and one glucokinase (Glk1). The relative amount of these kinases changes depending on a cell’s state and environment: Glk1 expression is induced in the absence of glucose and suppressed in high glucose (*9*) (Figure S1). We created a Glk1-GFP fusion expressed at the native locus and examined its behavior in different growth conditions. In the absence of glucose, Glk1-GFP was diffuse throughout the cytoplasm. When glucose-starved cells were exposed to glucose, Glk1-GFP assembled into large filament bundles. These filaments rapidly disassembled when glucose was removed (Figure 1A-C and movies S1-2). In contrast, Hxk1-GFP and Hxk2-GFP did not exhibit this behavior (Figure 1A). Glk1-GFP polymerization also occurred when starved cells were introduced to mannose or glucosamine, both substrates of Glk1 (Figure S2 and S3).

**Fig 1.**
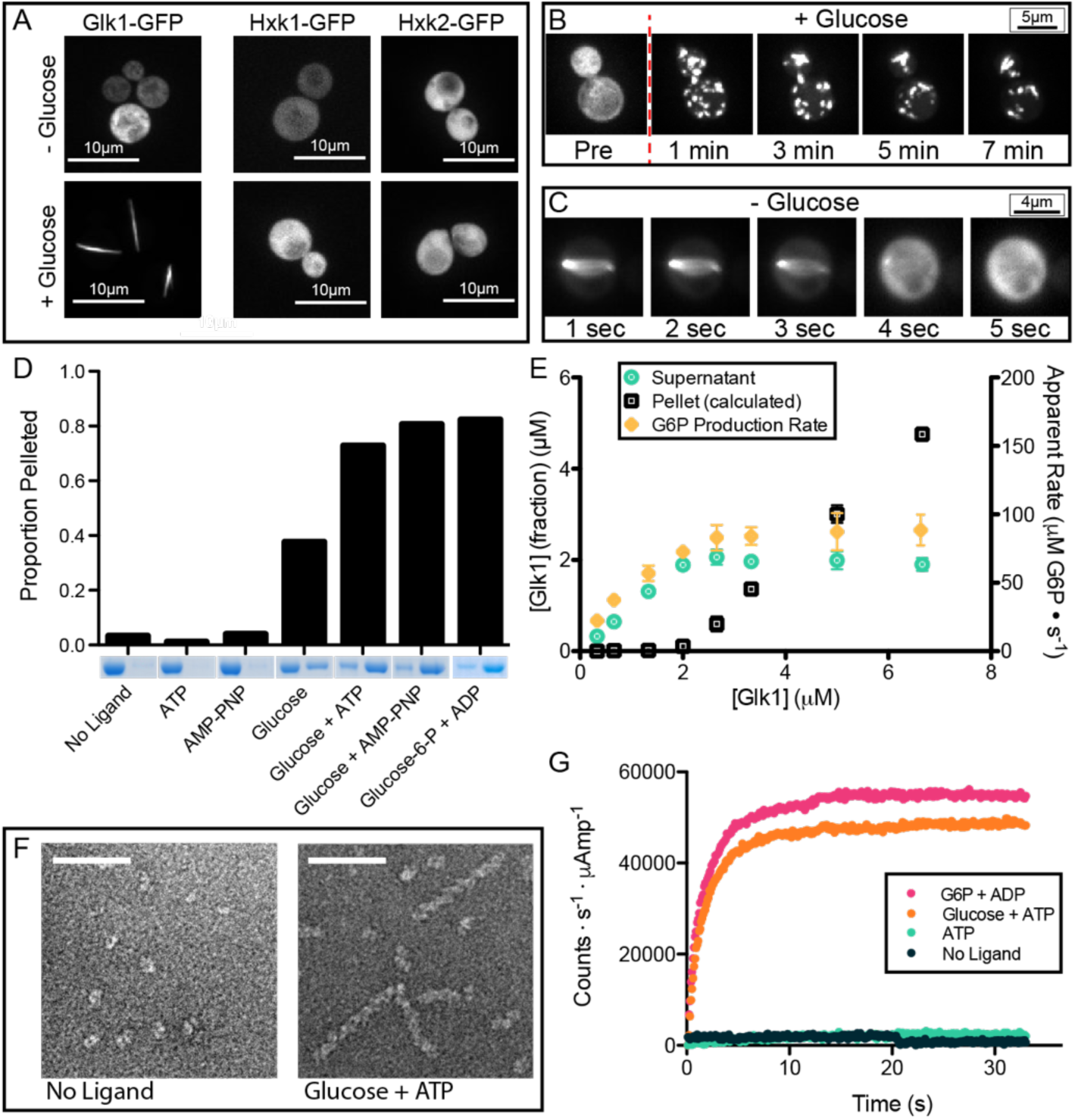
Glk1 forms filaments in response to its substrates at high enzyme concentration. **A)** Fluorescence images of stationary phase Glk1-GFP (*left*), Hxk1-GFP (*middle*), and Hxk2-GFP (*right*) cells before the addition of glucose (*top*) or after (*bottom*). Neither hexokinase forms higher-order structures in glucose; Glk1-GFP forms filaments. Scale bars = 10µm. **B, C)** Fluorescence images Glk1-GFP cells from a time-lapse. Glucose is added to the flow cell (*top*) or washed out (*bottom*). Glk1-GFP filaments form and dissolve rapidly in response to changes in the external glucose concentration. The filaments in B eventually coalesce into single filaments as in A and C. Scale bar is 5µm in B and 4µm in C. **D)** Purified Glk1 was ultracentrifuged with different ligand combinations. The supernatant (*left*) and pellet (*right*) of each condition were subjected to SDS-PAGE. Bar graph shows relative band intensities. Glk1 polymerizes in response to glucose. Polymerization is enhanced by ATP. Both substrate and product induce polymerization. **E)** Purified Glk1 concentration was varied in saturating glucose and ATP and assayed for enzyme activity (glucose-6-phosphate production rate) or ultracentrifuged. The concentration of Glk1 remaining in the supernatant was measured, and the amount in the pellet was inferred. **F)** Negative-stain electron micrographs of Glk1. 7.5µM Glk1 in the absence of ligands (*left*) or in saturating glucose and ATP (*right*), stained with uranyl formate. Scale bar is 50nm. **G)** 90-degree light scattering traces of Glk1 polymerization induced by rapid mixing with indicated substrates.

To understand what regulates Glk1 filament formation, we purified the enzyme and studied its polymerization *in vitro*. While a variety of other enzymes form condensates when cytosolic pH is decreased (*7*), purified Glk1 did not polymerize in response to pH changes (Figure S4). Rather, Glk1 polymerized in the presence of its enzymatic substrates (ATP and glucose, mannose, or glucosamine) or its products (ADP and sugar-6-phosphate). Modest polymerization occurred with N-acetylglucosamine and N-acetylglucosamine-6-phosphate, inhibitors of Glk1 activity that are neither substrates nor products (Figures 1D and S5) (*10*). Although Glk1 polymers form *in vivo* upon addition of fructose or galactose, these sugars do not induce Glk1 polymerization *in vitro*, suggesting the *in vivo* polymers assemble when the cell converts these sugars into glucose-6-phosphate (Figure S2 and S3) (*11, 12*).

Similar to other actin-fold protein polymers, Glk1 displays a critical concentration. Beneath 2µM Glk1 there was no polymerization. Above 2µM, the concentration of polymer increased with the total Glk1 concentration, while the concentration of unpolymerized Glk1 remained constant (Figure 1E). This is consistent with the lack of polymers in exponentially growing cells where Glk1 expression is suppressed by the presence of glucose. Indeed, Glk1-GFP polymers are observed when cells that express Glk1-GFP from a strong, constitutive promoter are grown in glucose (Figure S6). Consistent with the rapid polymerization observed *in vivo, in vitro* Glk1 polymerization was also rapid, reaching steady state in a matter of seconds following addition of ligands (Figure 1G).

### Glk1 polymerization inhibits glucokinase activity

Given that polymerization can either activate or inhibit enzyme activity (*6, 13*), we measured the rate at which Glk1 phosphorylated glucose as we varied Glk1’s concentration. Beneath the enzyme’s critical concentration (2µM), the rate of glucose-6-phosphate (G6P) production increased with Glk1 concentration. In contrast, above the critical concentration the rate of product formation was constant (Figure 1E). Thus, polymerization inhibits Glk1 activity, and, at any concentration above the critical concentration, the monomer-polymer equilibrium holds the monomer concentration constant, thereby keeping net enzymatic activity constant.

### Glk1 forms two stranded, anti-parallel helical filaments with novel inter-monomer contacts

We used negative stain electron microscopy to examine Glk1 oligomers. In the presence of substrates, Glk1 formed short, helical filaments (Figure 1F). The absence of larger aggregates suggests that the thick GFP-Glk1 filament bundles observed *in vivo* are driven by crowding (*14*), stabilization by filament binding proteins (*15, 16*), or higher-order structure formation driven by the dimerization of the GFP tag (*17*).

To try to understand why Glk1 polymerizes and Hxk1 and Hxk2 do not, we solved the crystal structure of Glk1 (PDB ID: 6P4X, Table S1). Comparing this structure to the *S. cerevisiae* Hxk2 structure (*18*) revealed interesting differences (Figure 2A). While the N-terminal helix is flush to the end of the domain in Hxk2, in Glk1 the N-terminal alpha helix extends beyond the large subdomain and contains a solvent-exposed phenylalanine (Figure S7A). In the hexokinases, the C-terminal alpha helix extends beyond the end of the small subdomain, but in Glk1 the C-terminal helix ends within the small subdomain, creating a hydrophobic patch (Figure S7B). Two loops are extended in Glk1 relative to the equivalent loops in Hxk2, spanning Glk1 residues 230-243 and 438-444 (Figure S7D).

**Fig 2.**
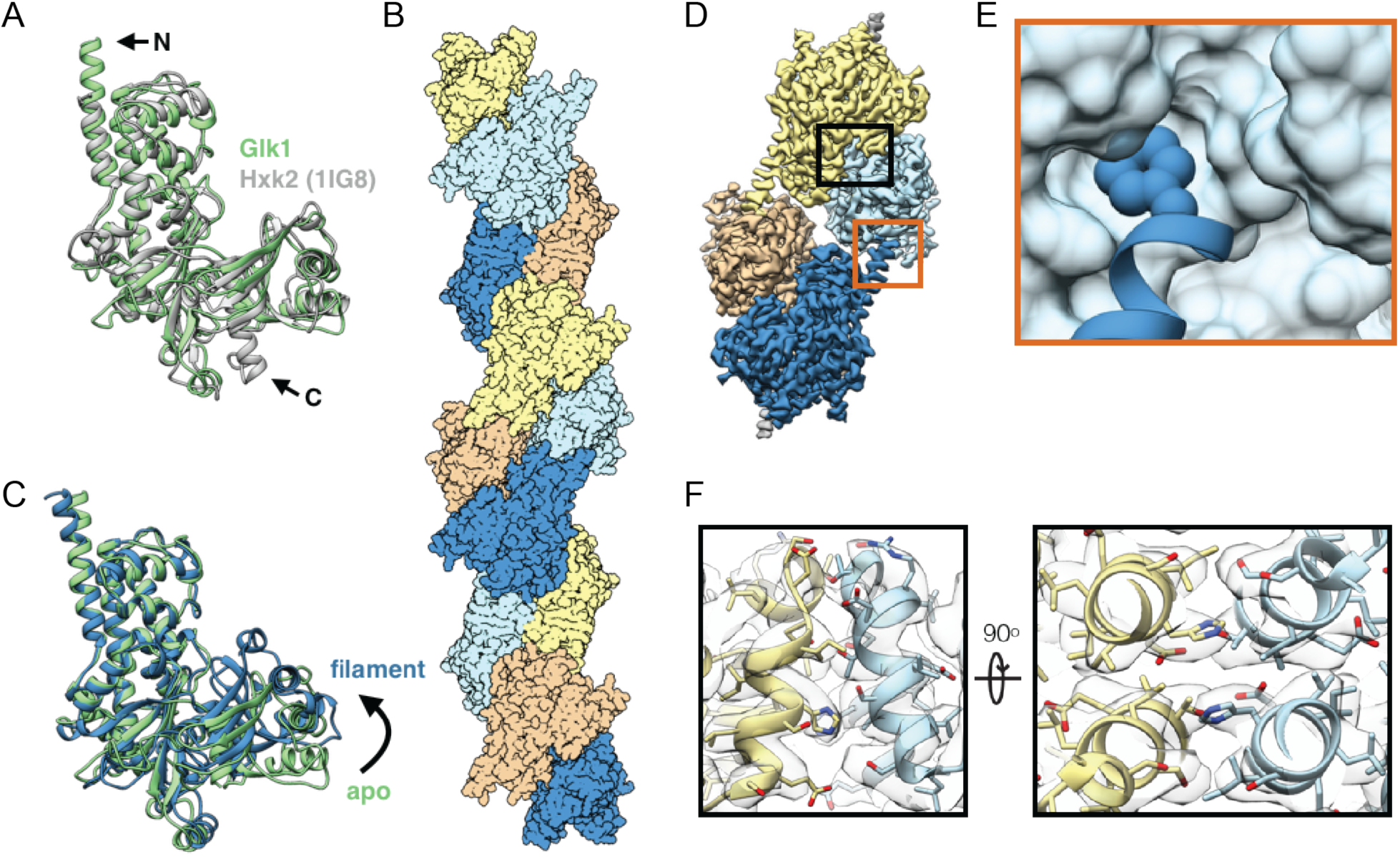
Glk1 forms anti-parallel, two-stranded filaments in its closed state. **A**: Superimposition of Glk1 crystal structure (*green*) with Hxk2 (PDB ID: 1IG8) (*18*) (*white*). The N-terminal helix of Glk1 extends further than Hxk2 (arrow, N), while the C-terminal helix of Hxk2 extends further (arrow, C). **B**: Surface representation of Glk1 filaments. Monomers along each strand are either orange/yellow or blue/cyan. **C**: Superimposition of the Glk1 crystal structure (*green*) with the Glk1 filament conformation (*blue*). The crystal structure is not ligand bound and is in the open state while the filament form is ligand bound and in the closed state. **D**: Electron density of four monomers in the Glk1 filament. A longitudinal contact is boxed in orange, and a lateral contact is boxed in black. **E**: Closeup of longitudinal contact with Phe3 represented as van der Wahls spheres and the next monomer represented as a surface. Phe3 of one monomer inserts into the hydrophobic pocket near the next monomers C-terminus. **F**: Two orthogonal close-ups of lateral filament contact. The helix-loop-helix from residue 371-393 of one monomer (*yellow*) binds the same region of the adjacent monomer (*blue*). Electron density is represented as transparent grey.

We used cryo-electron microscopy to determine the structure of Glk1 filaments (PDB ID: 6PDT, Table S2). This structure (3.8Å resolution) (Figure 2B) revealed that Glk1 forms two-stranded, antiparallel filaments. Monomers in the filament are in the closed state, consistent with Glk1 polymerizing when bound to ligand (Figure 2C) and with the other actin-like filaments (*4*). We also observed electron density consistent with glucose and ATP in the Glk1 active site, confirming that Glk1 filaments are ligand bound (Figure S8).

The interactions between Glk1 monomers along a strand of the filament differ from the conserved interactions seen in other polymer forming families in the Actin ATPase clan (Figure S9). In Glk1 filaments, the N-terminal, solvent-exposed Phe3 of one monomer inserts into the hydrophobic pocket at the C terminus of the next monomer in the same strand (Figure 2D-E). Lateral associations between strands are mediated by the helix-loop-helix between residues 371-393 interacting with the same region on the adjacent protein (Figure 2F). Furthermore, the interfaces between monomers along a single strand have contributions from loop 230-243. This loop is smaller in the hexokinases, which do not polymerize (Figure S7D-F).

### Glk1 polymerization evolved within the ascomycetes

The fungal glucokinases and hexokinases form separate clades that diverged at or before the origin of the Ascomycetes, one of the two major branches of the fungi. The group of yeast that contains *S. cerevisiae* (the Saccharomyceteceae) arose within the ascomycetes roughly 200 million years ago. The Glk1 homologs in most of Saccharomyceteceae contain four conserved motifs missing in both the other ascomycete Glk1 homologs and all ascomycete Hxk1/2 homologs. These motifs are in regions involved in Glk1 filament contacts: the N-terminus, loop 230-243, loop 438-444, and the C-terminus. To test if these motifs were predictive of polymerization, we purified several of the Glk1 homologs and Hxk1/2 homologs and tested their ability to polymerize. Of the enzymes tested, only Glk1 homologs that contained all four motifs polymerized in response to glucose and ATP (Figure S7 and S10). These results suggest that the ability of members of the Glk1 family to polymerize arose within the ascomycetes 150-200 million years ago, at the origin of the Saccharomyceteceae, and that one lineage within this group, the Kluveromycetes, has subsequently lost this ability (Figure 3A) (*19, 20*).

**Fig 3.**
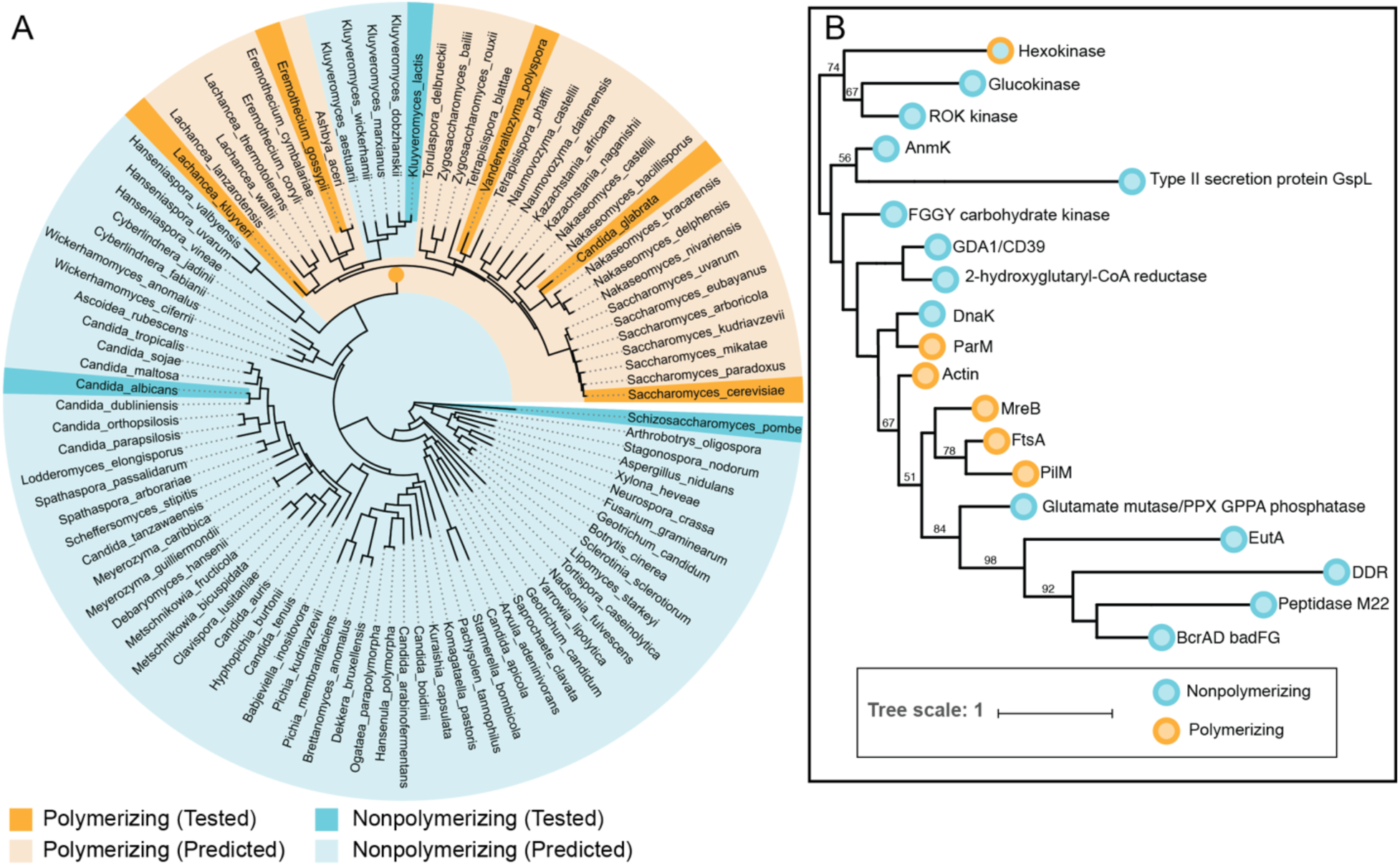
Glucokinase polymerization evolved independently of other actin-related polymers. **A)** Tree of ascomycetes as calculated by (*20*). Species whose Glk1 homologs polymerize are indicated in dark orange while those whose Glk1 homologs do not polymerize are marked in dark cyan. Species whose Glk1 homologs are predicted to polymerize based on their conservation of regions involved in intermonomer contacts (*pale orange*) and those predicted to not polymerize (*pale cyan*) are also indicated. The origin of Glk1 polymerization is indicated by the dark orange node. **B)** Phylogeny of Actin ATPase families. This summarizes phylogenetic analysis of 802 sequences from Actin ATPase protein families. A maximum likelihood tree was inferred under the LG+C20 substitution model in IQ-Tree (*28*). This displays the backbone structure of that ML tree with each protein family collapsed to display the relationships among families. Much of the backbone is uncertain; bootstrap supports are shown when >50. The tree suggests that the hexokinase family, which contains Glk1, forms a clade with ROK kinases and glucokinases, and is only distantly related to other actin families that are known to form polymers. Families that do not polymerize are cyan while families that do polymerize are orange. The full phylogeny is provided as Figure S11.

The protein families within the Actin ATPase clan previously reported to form polymers form a single clade. The protein family that contains Glk1 (the hexokinase family) is located outside of this clade, grouping with two other families of carbohydrate kinases: the glucokinase family and the ROK kinase family (Figure 3B and S11). This phylogenetic relationship is consistent with patterns of pairwise similarity observed between Hidden Markov Models of each family (Figure S12 and Table S3). Together, these results demonstrate that Glk1 polymerization evolved independently of other actin-fold polymers.

### Glk1 activity and polymerization are separable

Given that Glk1 polymerization has been conserved for millions of years, we examined how disrupting Glk1’s polymerization affected it’s enzymatic activity and the physiology of cells. To create a non-polymerizing Glk1 (NonPol-Glk1), we mutated the N-terminal phenylalanine involved in inter-monomer contacts to serine (Glk1-F3S). Pelleting assays confirmed that NonPol-Glk1 did not form polymers in response to glucose and ATP *in vitro* (figure 4A), nor did NonPol-Glk1-GFP form visible polymers in cells upon glucose addition (figure 4B). Assaying the rate of G6P production revealed NonPol-Glk1 was still enzymatically active but lacked the concentration-dependent inhibition observed in wild-type Glk1 above its critical concentration (figure 4C).

**Fig 4.**
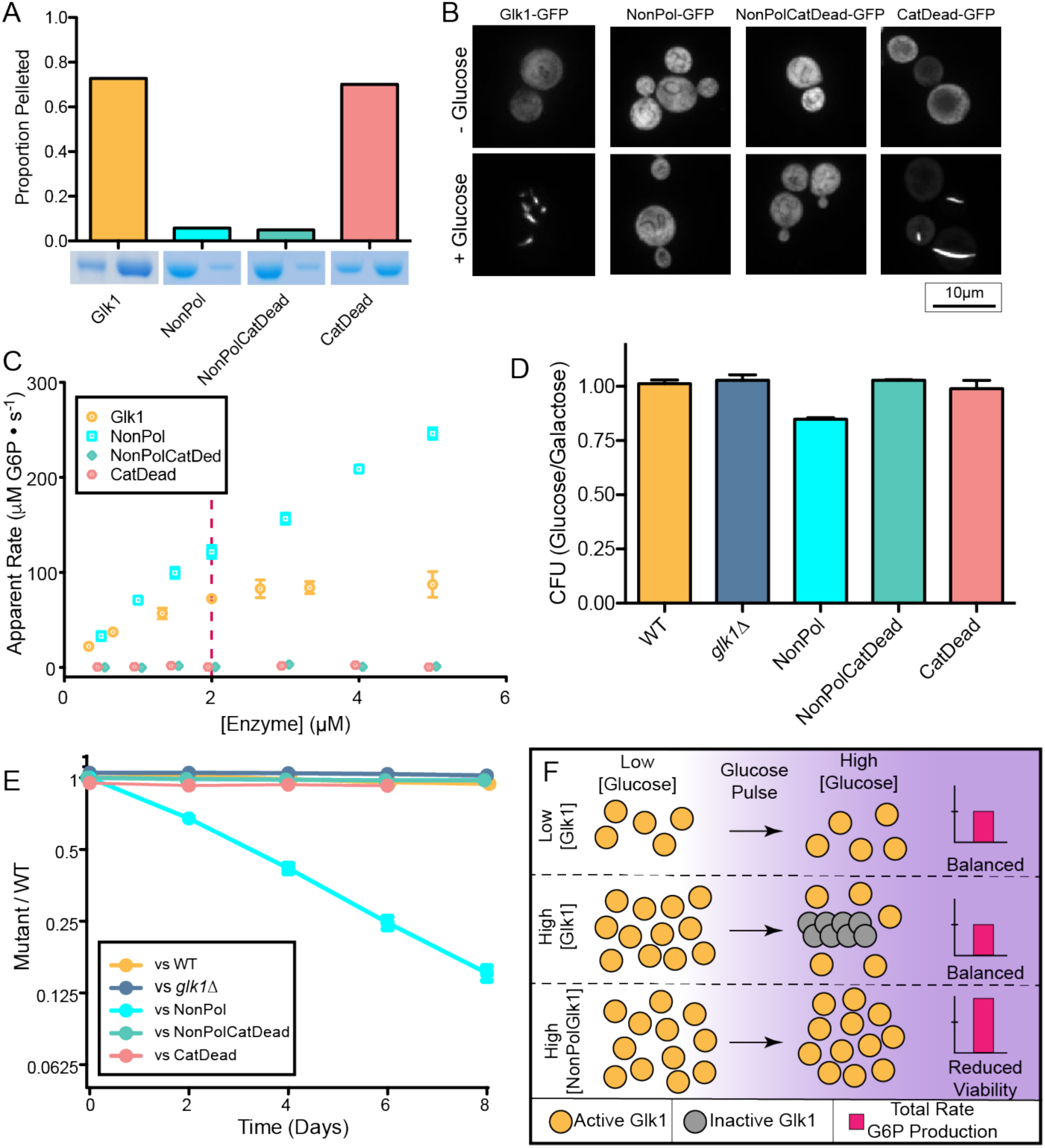
Glk1 polymerization inhibits Glk1 activity and elimination of polymerization causes fitness defects. **A)** 5µM purified non-polymerizing Glk1 (NonPol-Glk1), non-polymerizing catalytic-dead Glk1 (NonPolCatDead-Glk1), and catalytic-dead Glk1 (CatDead-Glk1) were ultracentrifuged with saturating glucose and ATP. The supernatant (*left*) and pellet (*right*) were subjected to SDS-PAGE. Bar graph shows relative band intensities. **B)** Fluorescence images of: Glk1-GFP (*left*), NonPol-Glk1-GFP (*middle-left*), NonPolCatDead-Glk1-GFP (*middle-right*), or CatDead-Glk1-GFP (*right*) stationary phase cells before (*top*) or after (*bottom*) refeeding glucose. Scale bar is 10µm. **C)** The rate of glucose-6-phosphate production at different concentrations of purified Glk1, NonPol-Glk1, NonPolCatDead-Glk1, and CatDead-Glk1. Glk1’s apparent rate does not increase beyond Glk1’s critical concentration (see Fig 1C). NonPol-Glk1 lacks concentration-dependent inhibition. NonPolCatDead-Glk1 and CatDead-Glk1 are inactive. **D)** The ratio of colonies grown on citrate buffered synthetic medium with glucose vs galactose after cells were conditioned in CBS-Galactose. NonPol-Glk1 cells die more frequently when refed glucose. **E)** mCherry labeled wild-type cells were competed against GFP labeled cells with different genotypes (wild-type, glk1?, NonPol-Glk1, NonPolCatDead-Glk1, CatDead-Glk1) through growth and dilution cycles in synthetic medium with glucose. The proportion of strains was measured after dilution by flow cytometry. **F)** Schematic of the effect of Glk1 polymerizations effect on glucokinase activity. When Glk1 concentration is high and glucose concentration increases, Glk1 polymerizes until the monomer concentration equals the critical concentration. Glk1 polymers lack enzyme activity: regardless of Glk1 concentration, the concentration of active enzyme is the same after glucose addition. When Glk1’s ability to polymerize is disrupted, it’s glucokinase activity is unconstrained, leading to fitness and viability defects.

To distinguish between the cellular effects of NonPol-Glk1’s lack of enzyme inhibition and the absence of Glk1 polymers, we mutated the catalytic lysine (K182A) (Figure S8B) (*21*) to create a catalytically dead Glk1 (CatDead-Glk1). We then combined these two mutations (F3S/K182A) to create non-polymerizing, catalytically dead Glk1 (NonPolCatDead-Glk1). As expected, CatDead-Glk1 formed polymers both in response to glucose in cells and to glucose and ATP *in vitro*, but did not generate G6P. NonPolCatDead-Glk1 neither formed polymer nor produced G6P (figure 4A-C).

### Non-polymerizing Glk1 reduces the fitness of yeast cells growing on sugars

When starved yeast are refed glucose, excess sugar kinase activity is toxic. This arises from an imbalance between the early steps of glycolysis, which consume ATP, and the late steps, which generate ATP (*22*). In yeast, Hxk1 and Hxk2 activity is inhibited by trehalose-6-phosphate, a metabolic intermediate that transiently accumulates as a result of elevated glucose-6-phosphate (*23*). Because Glk1 is not inhibited by trehalose-6-phosphate (*23*), we hypothesized that Glk1 polymerization prevents excessive Glk1 activity during the start-up of glycolysis when starved cells are refed glucose. Consistent with this model, when cells approaching stationary phase in galactose are refed glucose, 15% of cells containing NonPol-Glk1 die (figure 4D). This death is due to unregulated Glk1 activity and not lack of Glk1 polymers: cells that either lack Glk1, contain CatDead-Glk1, or contain NonPolCatDead-Glk1 showed no significant viability difference compared to wild-type. Together, these results suggest that Glk1 polymerization serves to limit the rate of glucose phosphorylation during the start-up of glycolysis.

Unregulated Glk1 activity is also detrimental to fitness over the entire growth cycle. We used differential fluorescent labeling to compare the fitness of wild-type cells against each of the mutants. We diluted the mixed cultures into fresh medium every 48 hours, measuring the proportion of each strain by flow cytometry. When growing on glucose, NonPol-Glk1 cells have a substantial fitness defect, averaging to a fitness cost of 6% over the entire 48-hour growth cycle (*24*). In contrast, none of the other Glk1 mutants or *glk1Δ* showed any impairment (figure 4E). Similar effects were observed when competing these strains on other sugars (figure S13).

## Discussion

This work demonstrates that the *S. cerevisiae* glucokinase, Glk1, polymerizes in response to its substrates and products. Polymerization inhibits Glk1 enzyme activity; only the monomeric pool is active. Thus, Glk1 polymerization governs the bulk rate of catalysis, with the Glk1 critical concentration setting the upper limit of flux through the entire Glk1 pool, regardless of enzyme or substrate concentrations (Figure 4F). This mode of self-regulation is not only robust to growth state or cell-to-cell variations in protein levels, it is reversible, allowing rapid adaptation to transient perturbations. In this sense, Glk1 polymerization behaves as a molecular surge protector, buffering the cell against not only nutrient spikes, but also the large differences in Glk1 concentrations that occur in different growth media. We argue that Glk1 polymerization evolved as a strategy to allow cells to adapt to environmental transitions occurring faster than the cell can change its protein levels. It is likely that other metabolic enzymes also polymerize in response to their substrates or products, and this may reflect a general mechanism cells use to rapidly adapt to stochastic changes in environmental conditions.

What pressures might have led to glucokinases evolving to polymerize? The group of yeast in which Glk1 polymerizes all lack the mitochondrial Respiratory Complex I, which reduces their total respiratory capacity (*25*). When glycolytic flux exceeds the rate of respiration it can lead to the buildup of metabolic byproducts and alter the redox balance within cells (*26, 27*). Thus, glucokinase polymerization may have evolved as an adaptation to cope with the imbalance between glycolytic flux and limited respiratory capacity during glucose pulses.

These findings reveal that polymerization can evolve in different ways from a given protein fold. The actin-like cytoskeletal polymers group into several distinct families, each forming a different filament ultrastructure. Phylogenetics and their conserved inter-monomer geometry (*5*) suggest that they share a common ancestor. In contrast, Glk1 polymerization arose independently of these other polymers (within the non-polymerizing glucokinases of the Ascomycetes) evolving a different geometry between monomers. Thus, it appears that polymerization has evolved in the same protein fold at least twice, each time selecting a different set of monomer-monomer interactions. The independent evolution of Glk1 polymerization, with its different inter-monomer interfaces, strengthens the hypothesis that the actin-like cytoskeletal polymers (actin, MreB, ParM, FtsA, MamK, and PilM) share a common polymeric ancestor. While this hypothesis was founded upon phylogenetic analysis (*4*), the ancient divergence of these families made it difficult to distinguish a single origin from several independent events. Our finding demonstrates that the conserved geometry seen in the actin-like cytoskeletal polymers is not an essential prerequisite for polymerization; instead, it likely reflects a common evolutionary origin.

## Acknowledgments

We thank Rachelle Gaudet for sharing beamtime and resources and for crystallographic advice, along with Christina Zimanyi, Lukas Bane, and Elizabeth May. We thank Jim Wilhelm for helpful discussions. We thank the Arnold and Mabel Beckman Cryo-EM Center at the University of Washington for access to electron microscopes. This work used NE-CAT beamlines (GM124165) at the APS (DE-AC02-06CH11357).

## Authors Contributions

*S. cerevisiae* strains were cloned by PRS and QAJ. Fluorescence microscopy was done by PRS and QAJ. PRS measured the relative expression of Glk1 by flow cytometry. Plasmid construction and protein purification, *in vitro* measurements of polymerization and enzymatic activity, Xray crystallography, Glk1 crystal structure refinement, and negative stain EM were done by PRS. PRS measured the viability of cells during glucose refeeding and measured the fitness of competed strains by flow cytometry. CryoEM sample preparation and data collection were performed by EML and AMD, cryoEM data was analyzed by EML and JMK, and the atomic model of Glk1 filaments was built by EML, DPF, and FD. Phylogenetic analysis of Actin ATPase families was performed by TAW. This project was conceived of by PRS, QAJ, ECG and AWM. PRS, EML, and TAW generated figures for this work and the paper was written by PRS, ECG, and AWM and edited by PRS, ECG, AWM, QAJ, EML, JMK, and TAW.

## Supplementary Materials

Materials and Methods

Figures S1-S13

Tables S1-S4

Movies S1-S2 available at http://garnerlab.fas.harvard.edu/Glk1/

References 29-59

