## Supplemental Information for "Independent evolution of polymerization in the Actin ATPase clan regulates hexokinase activity"

**This Supplement includes:**

Materials and Methods
Figs. S1 to S13
Tables S1 to S4
Captions for Movies S1 and S2
References (29-59)

**Other Supplementary Materials for this manuscript includes the following:**

Movies S1 and S2 available at http://garnerlab.fas.harvard.edu/Glk1

### Materials and Methods

##### Strains and Culture Conditions

Unless otherwise noted, *S. cerevisiae* was grown at 30˚C in CSM supplemented with the appropriate carbon sources. For viability/plating assays, yeast were grown in citrate buffered synthetic medium (CBS) (*21*) with appropriate carbon source.

##### Plasmid Construction and Cloning

Table S4 contains strains and plasmids used in this study. Constructs for the transformation of *S. cerevisiae* were generated by fusion PCR (*29*), and the cells were transformed with purified PCR product (*30*). Clones were verified by amplification of the genomic locus followed by Sanger sequencing of the purified product. For protein overexpression and purification from *E. coli,* glucokinases/hexokinases were cloned as $his_6$SUMO fusion constructs (*31*) into T7 plasmids by Gibson assembly (*32*). The plasmids were purified, and their sequences confirmed by Sanger sequencing using universal primers.

##### Media Transitions

Cells from cultures grown to saturation were loaded into a microfluidic cell (CellAsic), and their spent medium flowed over them. Cells were exchanged into fresh media containing 2% glucose. Media was pumped at 2PSI.

##### Microscopy

Images were taken on a Nikon Ti inverted microscope with a Yokogowa spinning disc confocal unit, 447, 488, 515, and 595 nm lasers, a Hamamatsu Orca camera operated with MetaMorph software. 8μm z-stacks were taken in 0.2μm increments. Z-stacks were converted to 8-bit, max intensity projected, and contrast adjusted in ImageJ (*33*).

The filament disassembly video (Movie S2) was taken using Nikon Ti-E inverted microscope (Nikon) equipped with a 60× objective (PlanApo, numerical aperture 1.4, oil), GFP filter (Chroma Technology), and a CoolSNAP charge-coupled device camera (Photometrics) using streaming acquisition of a single focal plane.

**Protein Purification**

BL21(DE3) Rosetta containing the $his_6$SUMO fusion plasmid were grown to $OD_{600}$ ~ 0.6 and induced overnight with 0.4mM IPTG at 16 ˚C. $His_6$SUMO fusion product was purified and cleaved as in (*31*). Proteins were further purified and exchanged into HKM-buffer (20mM HEPES-KOH pH 7.5, 100mM KCl, 1mM $MgCl_2$, 10% Glycerol (v/v), 10mM β-mercaptoethanol) using an S200 16/600 column on an AKTA FPLC. Fractions were pooled and concentrated. Aliquots were snap frozen in liquid nitrogen and stored at -80˚C until needed.

**Ultracentrifugation**

Unless otherwise noted, polymerization reactions were done in HKM-buffer and contained 5μM enzyme, 10mM Glucose, and 10mM MgATP. Polymerization reactions were spun in a TLA-100 (Beckman) at 100k rpm for 30 minutes at 30˚C. Supernatants were removed and added to equal volume of 2xSDS-Buffer. Pellets were resuspended by heating at 65˚C in 2 volumes of 1xSDS-Buffer. Fractions were subjected to SDS-PAGE, stained with Coomassie Blue G250, and band intensities were quantified in imageJ (*33*).

**Light Scattering**

Polymerization reactions were initiated by mixing 15µM Glk1 with an equal volume of 10mM Glucose and 10mM MgATP or 10mM Glucose-6-phosphate and 10mM MgADP using a SFA-20 rapid mixer (Hi-Tech Scientific), driven by a pneumatic drive unit (Hi-Tech Scientific). The 90-degree scattering of the solution at 315nm was measured using a Fluorolog-3 (Horiba).

**Crystallization, Diffraction, and Refinement**

Glk1 was exchanged into 5mM HEPES-KOH pH 7.5, 20mM KCl, 1mM $MgCl_2$, 0.5mM TCEP and concentrated to 11 mg/mL by centrifugal ultrafiltration (Amicon Ultra; Millipore). Conditions were screened around the conditions used in (*34*) as hanging drops. Final crystallization conditions were 2.4M $(NH_4)_2HPO_4$, 0.1M CHES pH 9.4 (2:1 protein:well-solution) with crystals forming after 5 days at 20˚C. Crystals were cryoprotected by briefly soaking in well solution supplemented with 18% glycerol before freezing.

X-ray data were collected at the Advance Photon Source beamline 24-ID-C at the Argonne National Laboratory. 180˚ of data was collected with a 10µm beam at 20% transmission. Reflection data were indexed, integrated, and scaled with HKL2000 (*35*). Initial phases were obtained using Phaser (*36*) for molecular replacement, using the *S. cerevisiae* hexokinase-2 structure (PBD ID: 1IG8) (*18*) as a search model. The model was built in Coot (*37*) and refined using Phenix.refine (*38*). Positional and B-factor refinement with TLS, torsion angle, and NCS restraints were used. Phosphates from the crystallization condition were modeled into densities of appropriate size that were coordinated by basic and polar residues. The asymmetric unit of the crystal contained six protein chains. Residues 51-59 were omitted from all chains because this loop had poor density. Besides this omission, chains A, C, and F contain residues 1-500, chains B and E contain residues 3-500, chain D contains residues 4-500. Chain A was used for analysis and generation of figures.

**CryoEM sample preparation and data collection**

Glk1 was buffer-exchanged into 20 mM K-HEPES pH 7.5, 100 mM KCl, 1mM $MgCl_2$, and 0.5 mM DTT using a desalting column. Glk1 filaments were then formed by incubating 20 μM Glk1 with 10 mM glucose, 10 mM $MgCl_2$, and 10 mM ATP for 20 minutes at room temperature. To prepare samples for cryoEM, 2.5 μl of protein was applied to glow-discharged CFLAT 2/2 holey-carbon grids (Protochips Inc.) and blotted away 4 times successively, before being plunged into liquid ethane using a Vitrobot (FEI co.). Data was collected on a Titan Krios (FEI co.) with a Quantum GIF energy filter (Gatan Inc.) operating in zero-loss mode with a 20 eV slit width. Movies were acquired on a K-2 Summit Direct Detect camera, operating in super-resolution mode with a pixel size of 0.525 Å/pixel, with 50 frames and a total dose of 90 electrons/$Å^2$. Leginon (*39*) software was used for automated data collection.

**CryoEM data processing**

Movie frames were aligned and dose-weighted using MotionCor2 (*40*), and CTF parameters were estimated using GCTF (*41*). Helices were picked manually from a subset of images using Relion (*42*), exported to cryoSPARC (*43*), and used to generate initial 2D classes which were then used as templates for picking from all micrographs in cryoSPARC. Further 2D classification, 3D refinement, and 3D classification was performed in cryoSPARC. Particles and parameters from cryoSPARC 3D refinement were then exported to Relion, and the final reconstruction was obtained using Relion 3D auto-refinement, with D1 and helical symmetry imposed. Helical symmetry parameters were also refined during Relion auto-refinement. Maps were sharpened using Relion post-processing, and resolution was estimated using the $FSC_{0.143}$ cutoff. Details of 3D reconstructions are summarized in Table S2.

**Atomic model refinement**

The Glk1 filament model was based on the solved crystal structure, PDB ID: 6P4X. First, small gaps in the model were resolved with RosettaCM (*44*) which used the model as a template and the cryoEM map EMD-20309 as a guide. Next, the model was further refined using the density and fragment sampling as described in (*45*). Finally, all-atom refinement was performed in Cartesian space utilizing the Rosetta FastRelax (*46*) protocol.

**Homolog Selection**

Glucokinase and hexokinase homologs for purification and polymerization testing were found by reciprocal BLAST (*47*) searches and were informed by synteny when possible. Protein alignments of the tested enzymes were made using Clustal Omega (*48*).

**Actin superfamily fold similarities**

Our analyses were based on a dataset of HMMs representing each member of the actin fold family obtained from the Pfam database (version 32.0). For families in which the N and C terminal domain halves were represented by different Pfam HMMs (hexokinase, glucokinase, and hydantoinase), we built new HMMs by searching the full-length query sequence of *S.*

*cerevisiae* Glk1 (UniProtKB – P17709) for the hexokinases, the *E. coli* Glk (UniProtKB – P0A6V8) for the glucokinases, and the *E. coli* HyuA (UniProtKB – Q46806) for the hydantoinases against the UniProtKB database using Jackhmmer (*49*). Pairwise similarity comparisons between the HMMs were performed using HHsearch from the HH-Suite 3 package (*50*).

**Actin fold phylogeny**

The phylogeny was based on a set of sequences assembled previously (*4*), but updated to include the full set of folds analyzed here. Representative sequences for each fold were selected by CD-HIT (*51*) clustering of the highly redundant set of sequences used to construct the corresponding HMMs. Sequences were aligned using the L-INS-i mode in mafft (*52*), poorly aligning regions of the alignment were identified and removed using the "gappyout" mode in trimAl (*53*), and the phylogeny was inferred under the LG+C20+F model in IQ-Tree 1.6.10 (*28*). Bootstrap supports were computed using the UFBoot2 algorithm (*54*).

**Glucokinase Activity Assays**

Glucose-6-phosphate production was measured in HKM-buffer at 30 ˚C. Timepoints were quenched in an equal volume of 100mM EDTA at 95 ˚C. Glucose-6-phosphate concentration was measured in each timepoint by a colorimetric enzyme-coupled reaction as in (*55*).

**Competition Experiments**

Strains were grown overnight in CSM with the appropriate carbon source, diluted into fresh medium and grown to mid exponential phase. mCherry labeled wild-type cells were mixed with

GFP labeled strains in equal numbers. Cells were back diluted 200-fold every 48 hours, and samples taken two hours later. The ratio of the two strains was measured by flow cytometry (Fortessa) and the data analyzed in FlowJo (Beckton and Dickinson).

**Viability Experiments**

Strains were conditioned in CBS/Galactose for 48 hours, being diluted to $1 \times 10^5$ cells/mL in fresh medium every 12 hours. Cells were harvested, washed twice in CBS with no carbon source, and plated onto either CBS/Glucose or CBS/Galactose. Colonies were counted after three days at 30˚C.

### Supplemental Figures

**Figure S1 – Glk1 expression under different growth conditions.**

Green fluorescence of cells expressing Glk1-GFP from the Glk1 locus were measured by flow cytometry while growing in glucose or 12 hours after glucose was depleted from the culture (*no Glucose*). Middle line in each violin represents the median of the distribution. Upper and lower lines represent top and bottom quartiles. Glk1 expression increases in the absence of glucose.

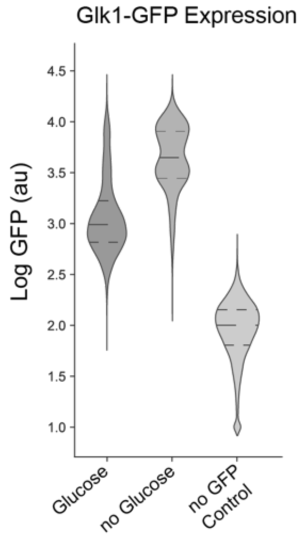


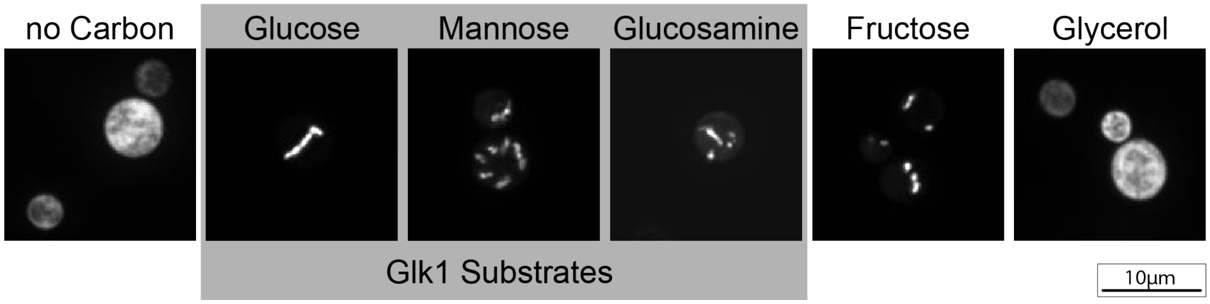


**Figure S2 - Glk1-GFP polymerizes in cells in response to several sugars.**

Maximum intensity projections of confocal z-stacks of Glk1-GFP cells. Cells were grown to saturation and washed in media containing no carbon. Cells were refed the sugar indicated, and fluorescence micrographs were taken 1 minute after refeeding. Carbon sources that are Glk1 substrates are boxed in gray. Scale bar is 10µm.

**Figure S3 – Galactose triggers Glk1-GFP polymerization only in galactose conditioned cells.**

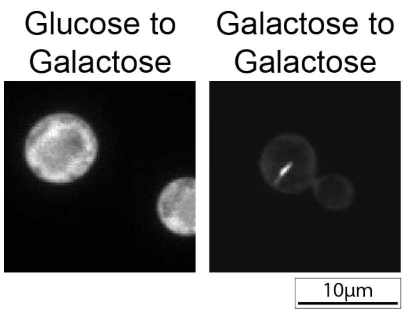


Fluorescence micrographs of cells expressing Glk1-GFP. Cells were either grown to saturation in glucose containing medium or galactose containing medium. Glk1-GFP polymerizes in galactose conditioned cells when refed galactose but not in glucose conditioned cells. This is likely due to the suppression of the galactose transporter (Gal2) and galactokinase (Gal1) by the presence of glucose (*56*). Scale bar is 10 μm.

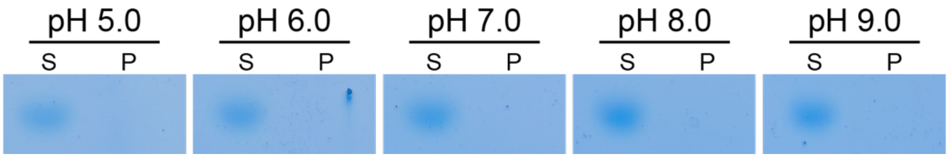


**Figure S4 - Glk1 does not polymerize in response to change in pH.**

10μM Glk1 was ultracentrifuged at pH values from 5.0 to 9.0, and the supernatant (S, *left*) and pellet (P, *right*) were subjected to SDS-PAGE and stained with Coomassie blue.

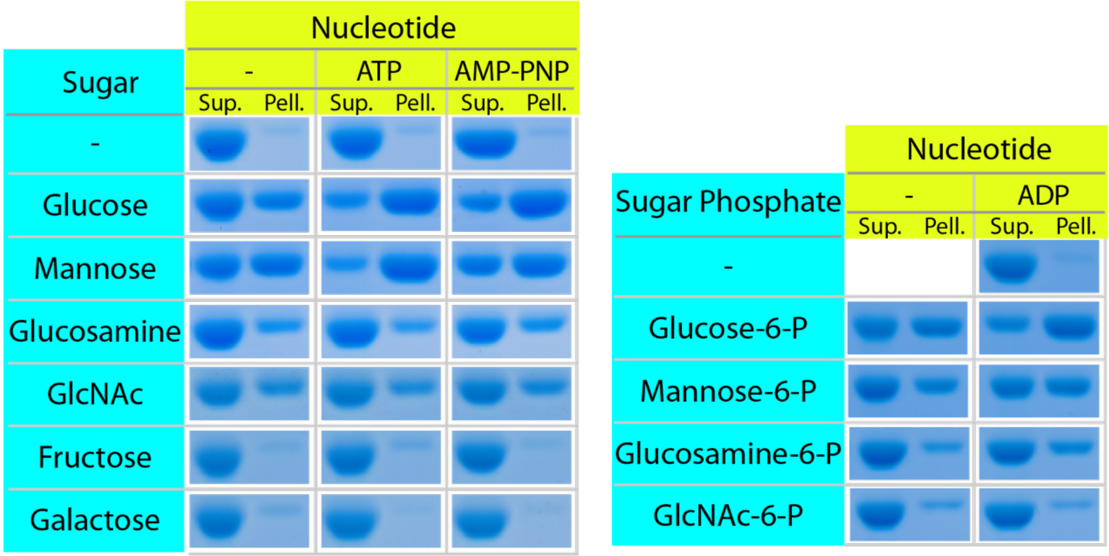


**Figure S5 – Glk1 polymerization in the presence of different ligands.**

Purified Glk1 was ultracentrifuged in the presence of different ligand combinations. The supernatant (*left*) and pellet (*right*) of each condition were subjected to SDS-PAGE. Glk1 polymerizes in response to its substrates (glucose, mannose, and glucosamine), inhibitors (GlcNAc and GlcNac-6-P), and products (glucose-6-P, mannose-6-P, and glucosamine-6-P), but not to other sugars (fructose and galactose).

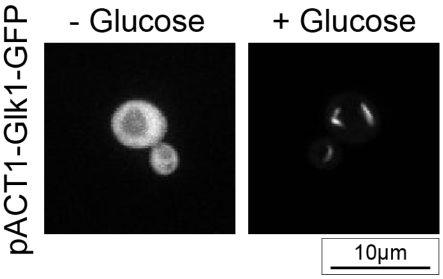


**Figure S6 - Fluorescence micrographs of cells expressing Glk1-GFP under the Actin promoter.**

Cells were harvested growing exponentially on glucose and imaged in the absence of glucose (*left*) or in the presence of glucose (*right*). Strong, constitutive expression of Glk1-GFP divorces cell state from polymer presence. Scale bar is 10µm.

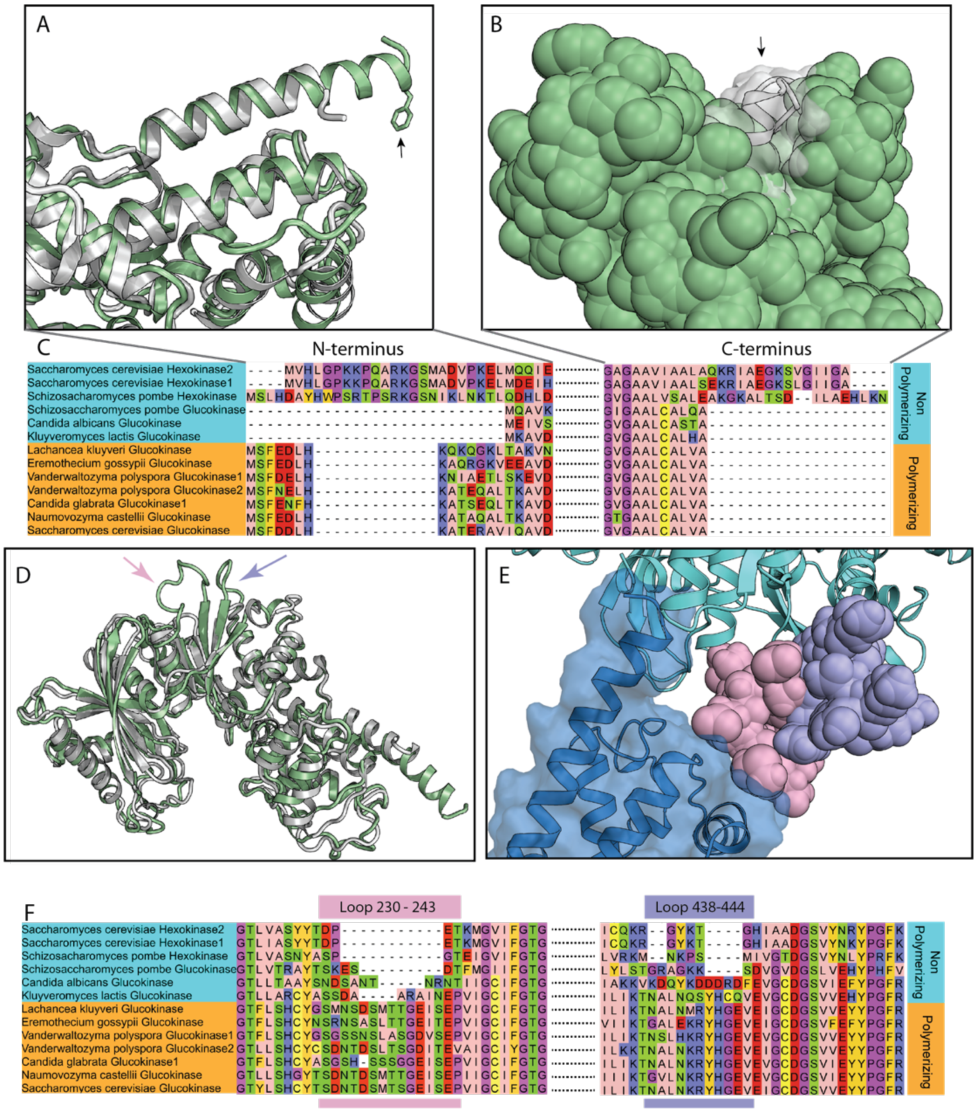


**Figure S7 - Residues involved in Glk1 filament contacts are conserved amongst polymerizing glucokinases but not amongst non-polymerizing glucokinases or hexokinases.** **A)** Comparison of the N-terminus of Hxk2 (*white*, PDB ID: 1IG8) and Glk1 (*green*). The N-terminal helix of Glk1 extends beyond the body of the protein, while the N-terminal helix of Hxk2 ends flush to the body. Glk1's helix contains a solvent exposed phenylalanine. **B)** The C-terminal helix of Hxk2 (*white ribbon/transparent surface*) extends beyond that of Glk1 (*green spheres*). This means that Hxk2 does not have the hydrophobic pocket involved in Glk1 polymerization. **C)** Alignment of all tested enzymes. The N terminus (MSF(e/d)(e/d)LHK) and

C-terminus (LCALVA) are conserved amongst enzymes that polymerize and divergent amongst enzymes that do not. **D)** The loop from Glk1 residue 230-243 (*pink arrow*), and Glk1 residue 438-444 (*purple arrow*) are extended relative to their corresponding loops in Hxk2 (*white*). **E)** Close up of longitudinal interface between two monomers in a Glk1 filament. Loop 230-243 (*pink spheres*) contacts the next monomer (*blue ribbon and surface*), and loop 438-444 (*purple spheres*) packs tightly against loop 230-243. **F)** Alignment of all tested enzymes in loop regions. Both loops are extended in all polymerizing enzymes and are truncated or divergent in non-polymerizing enzymes.

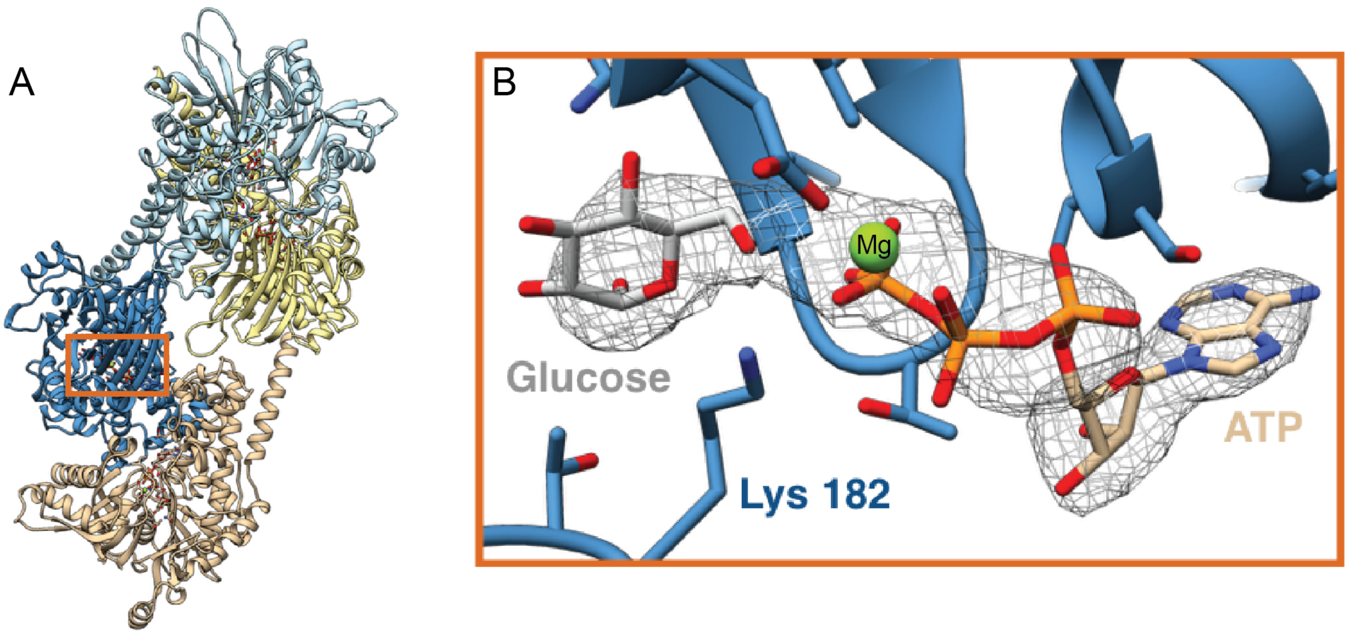


**Figure S8 - Glk1 filaments have magnesium and glucose and ATP or G6P and ADP bound in their active site.**

**A)** Cartoon representation of four monomers within a Glk1 filament showing where Glucose and ATP are modeled within the filament. The orange box indicates the location of the ligands shown in B. **B)** Ligand density (*gray mesh*) is present in the cryoEM Glk1 filament electron density. Glucose and ATP vs G6P and ADP cannot be distinguished at this resolution. Here we have modeled glucose and ATP. The catalytic lysine (Lys182) can be seen coordinating the -OH on the 6-carbon of glucose and the -OH on the gamma-phosphate of ATP.

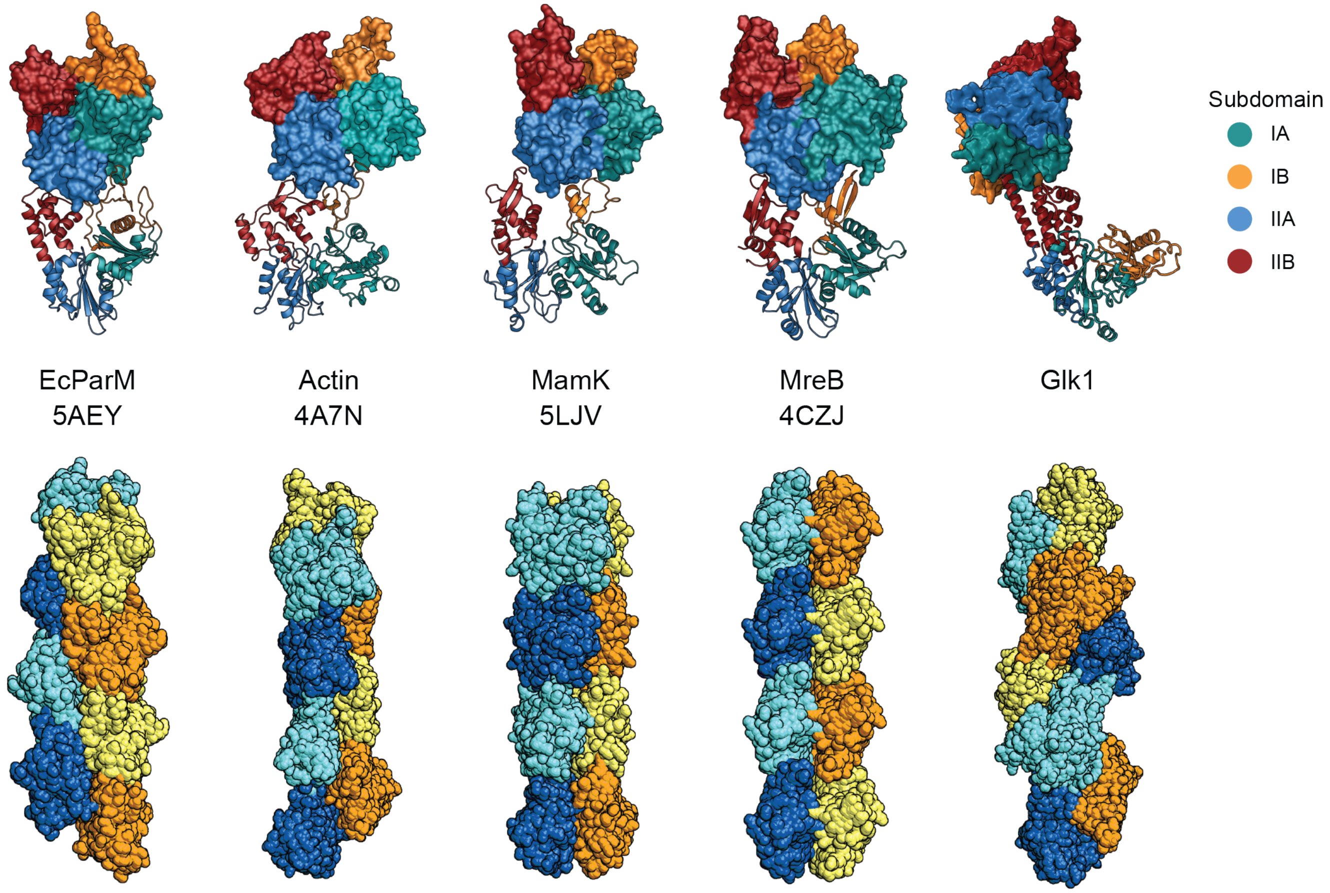


**Figure S9 - Intermonomer geometry conserved in other actin-related filaments is not present in Glk1 filaments.**

*Top*: Amongst all other actin-related filaments, along a single strand, subdomain IB and IIB contact subdomain IA and IIA respectively. In Glk1 filaments, subdomain IIB contacts subdomain IA (*5*, *57-59*). The top monomer is represented as a surface while the bottom monomer is represented as a ribbon.

*Bottom*: Despite the conserved geometry along strands in the cytoskeletal polymers, filaments are still able to achieve a variety of structures through varying monomer shape and lateral interactions

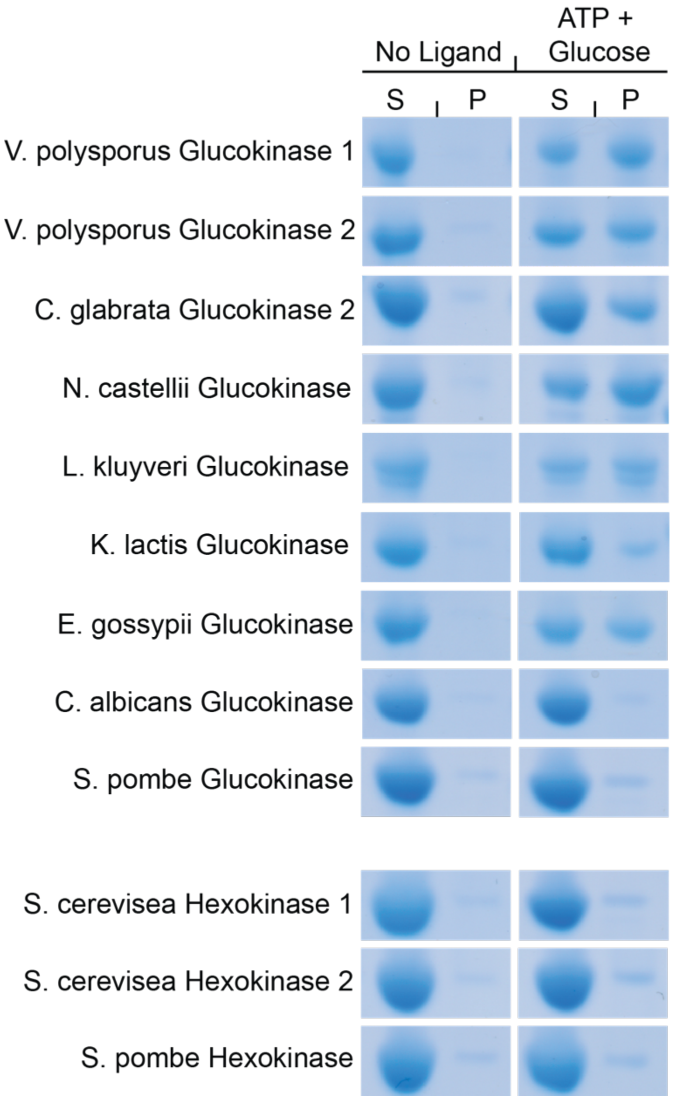


**Figure S10 - Polymerization of Glk1 homologs.**

Purified Glk1 homologs (glucokinases) and Hxk1/2 homologs (hexokinases) were ultracentrifuged in the absence of ligand (*left*) or the presence of glucose and ATP (*right*). The supernatant (*S, left*) and pellet (*P, right*) were subjected to SDS-PAGE and stained with Coomassie blue. Some other glucokinases polymerize in the presence of Glucose and ATP while hexokinases do not.

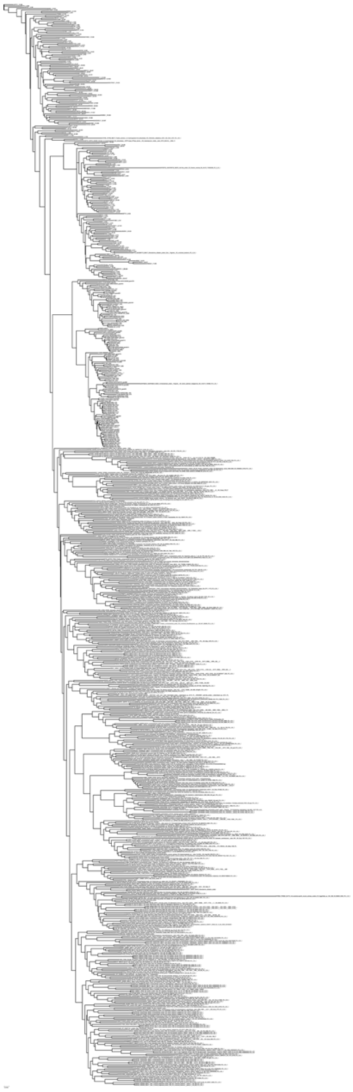

**Figure S11 - Maximum likelihood tree of the actin fold**.

A maximum likelihood tree of 802 actin fold proteins inferred under the LG+C20 substitution model in IQ-Tree (*28*). A high-resolution version of this image is available in the image files.

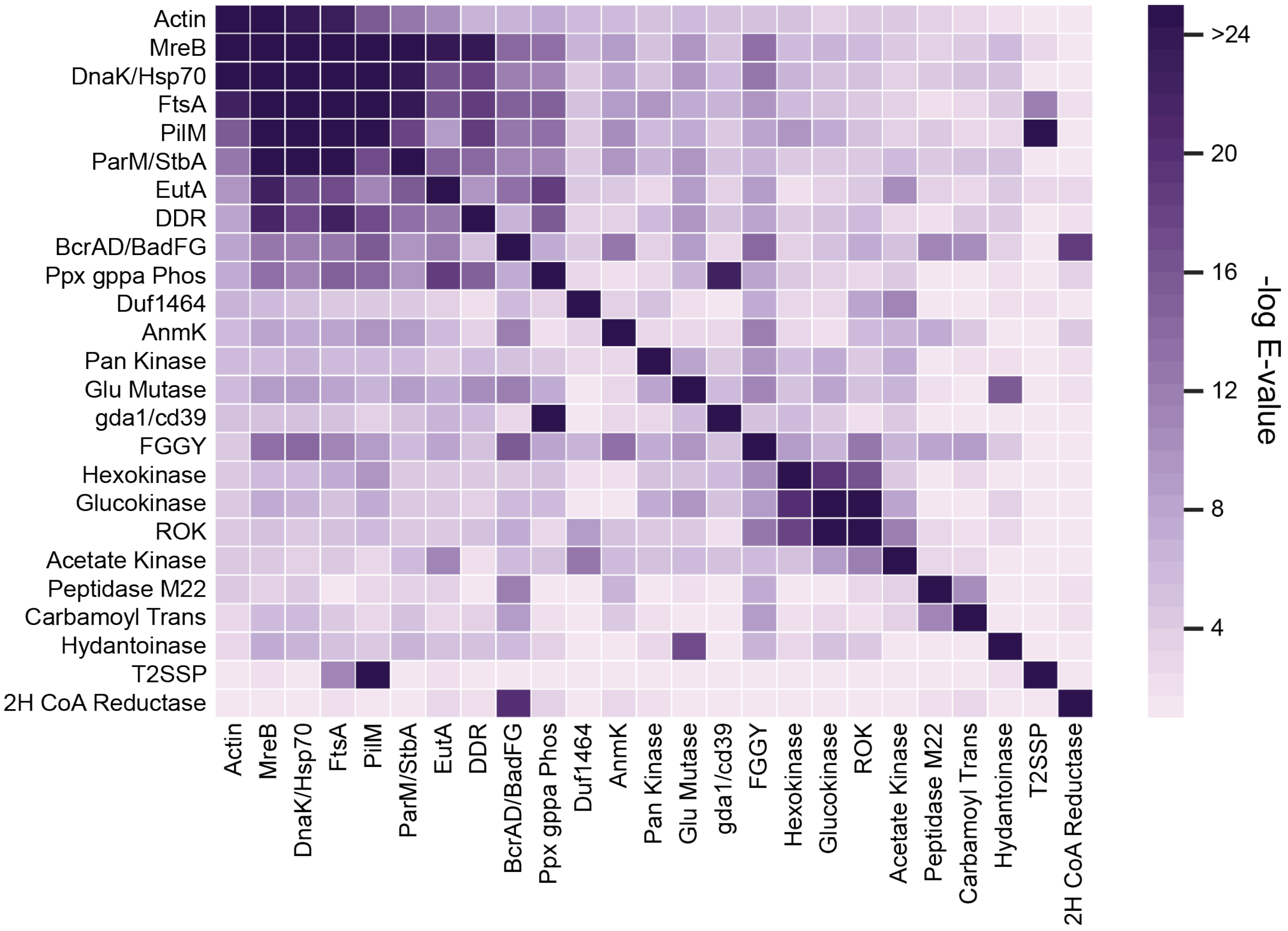


**Figure S12 - Heatmap representing the e-value for the comparison of HMM for each protein family in the Actin ATPase clan.**

The values are reported as negative logs of the e-value. Note that Glk1 is in the hexokinase, not the glucokinase family in this classification.

**Figure S13 - NonPol-Glk1 cells have growth defects in a variety of sugars, including galactose, which is not its substrate.** Wild-type W303 labeled with mCherry were competed against green labeled W303 (wild-type, glk1Δ, NonPol-Glk1, NonPolCatDead-Glk1, CatDead-Glk1) through repeated growth and dilution cycles in CSM-Mannose (*top*), CSM-Fructose (*middle*), or CSM-Galactose (*bottom*). The relative proportion of the strains were measured after each dilution by flow cytometry.

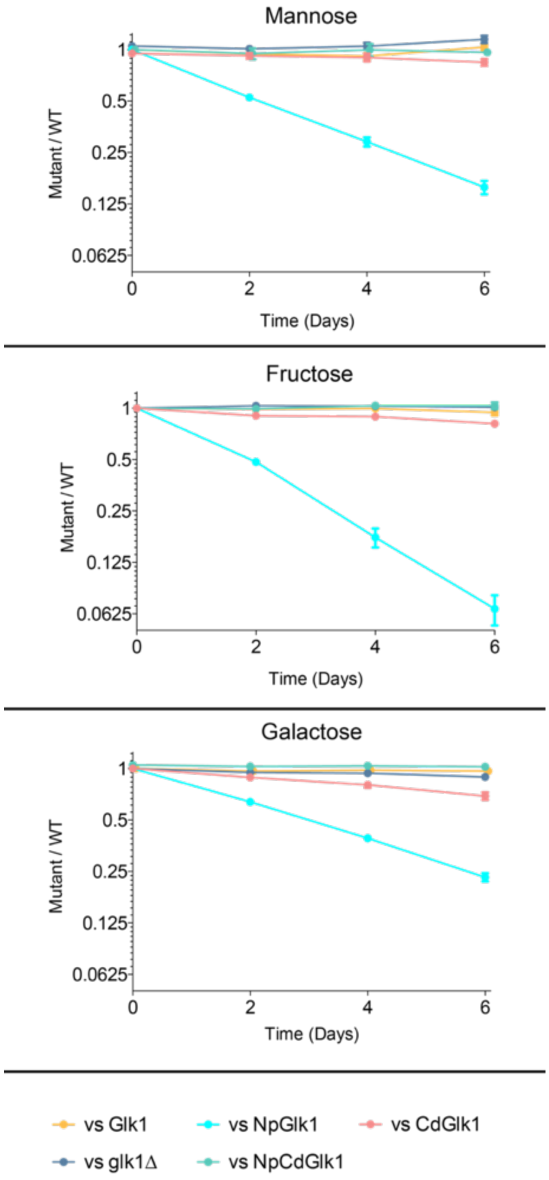

### Supplemental Tables

##### Table S1: Crystallography Statistics

| | Glk1 (PDB ID: 6P4X) |
|---|---|
| **Data Collection** | |
| Wavelength (Å) | 0.9793 |
| Resolution Range (Å) | 155.8 - 3.59 (3.718 - 3.59) |
| Space Group | P 43 21 2 |
| Unit Cell (a,b,c) | 174.5, 174.5, 323.947 |
| Unit Cell (α,β,γ) | 90, 90, 90 |
| Number of Crystals | 1 |
| Total Reflections | 122662 (12052) |
| Unique Reflections | 61332 (6026) |
| Redundancy | 2.0 (2.0) |
| Completeness (%) | 99.97 (99.95) |
| Mean I/σ (I) | 8.28 (1.56) |
| R-merge | 0.1003 (0.4831) |
| R-meas | 0.1418 (0.6832) |
| R-pim | 0.1003 (0.4831) |
| CC 1/2 | 0.988 (0.638) |
| **Refinement** | |
| Resolution Range (Å) | 155.8 - 3.59 (3.718 - 3.59) |
| R-work | 0.2652 (0.3358) |
| R-free | 0.3001 (0.3565) |
| Number of Atoms | 22953 |
| Protein | 22888 |
| Ligands | 65 |
| Protein Residues | 2944 |
| Ramachandran Plot | |
| Favored (%) | 98.73 |
| Allowed (%) | 1.27 |
| Outliers (%) | 0.0 |
| RMS (bonds) | 0.004 |
| RMS (angles) | 1.47 |
| Average B-factor | 89.65 |
| Protein | 89.69 |
| Ligands | 76.21 |
| Number of TLS Groups | 20 |

#### Table S2: CryoEM Statistics

| | **Glk1 filament** (EMD-20309, PDB 6PDT) |
|---|---|
| **Data collection** | |
| Electron microscope | Titan Krios |
| Voltage (kV) | 300 |
| Electron detector | K2 summit |
| Electron dose ($e^-$/Å$^2$) | 90 |
| Pixel size (Å) | 1.05 |
| **Reconstruction** | |
| Point group symmetry | D1 |
| Refine helical symmetry | 120.4°, 60.1Å |
| Particles | 56,778 |
| Resolution (0.143 fsc) (Å) | 3.8 |
| **Model composition** | |
| Protein residues | 500 |
| Ligands | ATP, glucose, Mg |
| **Validation** | |
| Clashscore | 1 |
| Poor rotamers (%) | 0 |
| **Ramachandran plot** | |
| Favored (%) | 98 |
| Allowed (%) | 2 |
| Outliers (%) | 0 |

**Table S3:** Table of negative log e-values for the pairwise comparison of the HMM for each protein family in the Actin ATPase clan. A high-resolution version of this table is in the image files.

| | Actin | MreB | DnaK/Hsp70 | FtsA | PilM | ParM/StbA | EutA | DDR | BcrAD/BadFG | ppx_gppa | DUF1464 | AnmK | Pan_Kinase | Glu_Mutase | gda1/cd39 | FGGY | Hexokinase | Glucokinase | ROK | Ace Kinase | Pep_M22 | Carbam_Trans | Hydant | T2SSP | 2H_CoA_red |
|---|---|---|---|---|---|---|---|---|---|---|---|---|---|---|---|---|---|---|---|---|---|---|---|---|---|
| Actin | 171.333 | 34.443 | 23.917 | 23.026 | 15.53 | 12.455 | 10.637 | 6.908 | 6.502 | 7.543 | 5.809 | 5.915 | 5.006 | 5.599 | 4.768 | 5.185 | 4.962 | 3.912 | 4.605 | 4.51 | 3.507 | 2.847 | 2.04 | 1.609 | 1.47 |
| MreB | 36.311 | 142.018 | 87.162 | 50.251 | 35.36 | 29.101 | 23.719 | 24.197 | 13.826 | 13.553 | 6.571 | 8.623 | 5.404 | 9.409 | 5.133 | 13.285 | 5.843 | 6.502 | 5.521 | 4.423 | 3.612 | 4.744 | 6.119 | 2.765 | 1.309 |
| DnaK/Hsp70 | 24.798 | 83.67 | 213.878 | 41.184 | 28.452 | 23.699 | 16.68 | 17.728 | 12.206 | 11.176 | 4.075 | 7.849 | 5.279 | 9.283 | 5.776 | 12.94 | 6.32 | 4.816 | 5.404 | 3.352 | 4.423 | 5.36 | 5.021 | 1.427 | 1.139 |
| FtsA | 22.284 | 47.399 | 40.859 | 160.018 | 97.065 | 23.719 | 16.149 | 18.722 | 14.636 | 14.732 | 4.804 | 8.68 | 9.316 | 7.094 | 6.32 | 9.657 | 5.915 | 5.279 | 4.51 | 3.73 | 1.833 | 2.976 | 4.075 | 11.596 | 2.04 |
| PilM | 15.936 | 36.008 | 29.671 | 100.783 | 146.724 | 17.728 | 9.21 | 18.441 | 12.86 | 13.72 | 4.678 | 10.127 | 6.119 | 7.156 | 4.343 | 8.422 | 9.35 | 7.07 | 5.221 | 3.507 | 4.343 | 2.551 | 3.244 | 27.867 | 1.386 |
| ParM/StbA | 12.717 | 27.946 | 24.264 | 25.233 | 16.873 | 133.288 | 14.592 | 13.816 | 10.925 | 11.107 | 4.343 | 9.361 | 6.32 | 9.272 | 5.15 | 6.97 | 4.269 | 4.343 | 4.51 | 6.075 | 4.722 | 4.791 | 5.279 | 0.236 | 1.386 |
| EutA | 9.987 | 22.62 | 16.118 | 17.391 | 11.251 | 15.33 | 389.137 | 9.327 | 13.41 | 18.722 | 4.269 | 4.615 | 2.604 | 9.21 | 3.817 | 8.805 | 2.303 | 3.381 | 4.343 | 10.232 | 3.863 | 2.976 | 4.51 | 2.513 | 2.957 |
| DDR | 8.335 | 21.894 | 17.322 | 22.237 | 17.356 | 13.228 | 12.94 | 583.247 | 6.502 | 15.936 | 3.54 | 3.612 | 6.166 | 9.721 | 5.319 | 7.775 | 4.343 | 5.051 | 5.599 | 2.526 | 2.04 | 4.343 | 4.605 | 0.821 | 2.04 |
| BcrAD/BadFG | 7.799 | 12.983 | 11.884 | 12.94 | 15.713 | 9.485 | 11.959 | 5.279 | 296.34 | 7.236 | 4.678 | 12.429 | 3.963 | 8.874 | 3.016 | 14.201 | 3.689 | 5.339 | 7.506 | 5.221 | 10.871 | 10.483 | 3.817 | 0.511 | 18.777 |
| ppx_gppa | 7.562 | 13.074 | 10.925 | 14.614 | 14.002 | 9.485 | 18.965 | 14.509 | 7.339 | 260.885 | 2.996 | 2.04 | 2.501 | 6.725 | 22.62 | 7.849 | 4.343 | 3.912 | 2.919 | 4.2 | 1.05 | 1.661 | 2.207 | 1.347 | 3.507 |
| DUF1464 | 6.266 | 5.843 | 4.934 | 4.605 | 4.135 | 4.656 | 3.507 | 1.897 | 5.547 | 3.772 | 296.34 | 3.817 | 5.185 | 2.207 | 0.4 | 7.293 | 2.781 | 2.674 | 8.079 | 11.418 | 0.968 | 1.661 | 2.303 | 1.897 | 1.386 |
| AnmK | 6.032 | 8.079 | 7.354 | 8.468 | 9.641 | 9.115 | 5.714 | 3.324 | 12.148 | 2.207 | 3.863 | 291.33 | 3.147 | 2.749 | 2.882 | 11.652 | 2.919 | 2.303 | 5.655 | 6.812 | 7.581 | 4.343 | 0.315 | 1.079 | 4.2 |
| Pan_Kinase | 5.684 | 5.655 | 6.377 | 5.952 | 5.745 | 5.745 | 4.605 | 5.714 | 5.051 | 4.135 | 2.847 | 2.996 | 156.863 | 8.335 | 4.343 | 9.657 | 5.684 | 7.118 | 4.017 | 7.25 | 1.204 | 1.897 | 1.772 | 0.598 | 2.12 |
| Glu_Mutase | 5.627 | 9.028 | 8.948 | 8.079 | 6.438 | 8.517 | 7.195 | 10.009 | 11.564 | 7.024 | 1.661 | 2.718 | 8.146 | 313.152 | 5.714 | 11.418 | 5.36 | 7.775 | 5.036 | 6.571 | 1.833 | 0.942 | 15.713 | 0.821 | 1.833 |
| gda1/cd39 | 4.962 | 4.867 | 4.841 | 4.867 | 3.772 | 5.15 | 6.98 | 5.521 | 3.058 | 24.335 | 1.139 | 2.83 | 2.538 | 5.627 | 269.913 | 5.36 | 6.075 | 4.135 | 2.207 | 4.269 | 0.545 | 0.462 | 0.329 | 1.609 | 1.022 |
| FGGY | 4.744 | 13.553 | 13.978 | 11.043 | 9.21 | 6.075 | 8.377 | 5.473 | 15.936 | 8.294 | 6.571 | 13.174 | 7.524 | 9.987 | 5.167 | 211.368 | 8.805 | 6.571 | 12.289 | 6.725 | 8.079 | 8.874 | 4.343 | 0.274 | 1.022 |
| Hexokinase | 4.636 | 6.166 | 6.075 | 7.024 | 9.79 | 4.343 | 4.2 | 4.2 | 4.135 | 5.449 | 2.364 | 2.12 | 5.279 | 5.382 | 5.991 | 10.724 | 282.119 | 19.499 | 16.281 | 4.269 | 0.654 | 2.847 | 1.661 | 0.598 | 1.05 |
| Glucokinase | 4.605 | 7.002 | 6.571 | 5.426 | 7.024 | 4.733 | 4.135 | 3.863 | 6.075 | 5.521 | 1.609 | 1.609 | 7.013 | 9.641 | 5.24 | 8.948 | 20.318 | 208.254 | 30.29 | 7.929 | 0.734 | 1.609 | 3.963 | 0.916 | 0.713 |
| ROK | 4.51 | 5.298 | 4.2 | 5.259 | 5.684 | 4.135 | 4.017 | 5.051 | 7.621 | 2.937 | 8.874 | 5.497 | 4.269 | 4.423 | 1.833 | 12.717 | 17.951 | 29.995 | 165.45 | 11.675 | 2.631 | 1.966 | 2.865 | 1.05 | 1.609 |
| Acetate Kinase | 4.343 | 4.51 | 3.612 | 4.017 | 3.101 | 5.655 | 10.82 | 2.04 | 5.809 | 5.426 | 12.405 | 5.521 | 5.167 | 6.166 | 5.067 | 6.075 | 4.934 | 8.568 | 11.761 | 291.042 | 2.703 | 3.244 | 0.261 | 1.109 | 1.079 |
| Pep_M22 | 4.017 | 3.65 | 4.2 | 1.079 | 3.016 | 3.912 | 3.324 | 1.609 | 11.533 | 1.427 | 1.204 | 6.377 | 1.715 | 1.833 | 0.58 | 7.419 | 1.171 | 0.58 | 2.996 | 3.507 | 219.522 | 10.127 | 0.223 | 1.204 | 1.897 |
| Carbamoyl_Trans | 3.058 | 5.878 | 5.809 | 4.269 | 2.9 | 4.768 | 3.037 | 3.73 | 8.948 | 1.772 | 1.427 | 4.605 | 2.453 | 1.427 | 0.371 | 8.74 | 2.354 | 2.957 | 1.897 | 3.442 | 10.982 | 334.791 | 0.562 | 0.994 | 2.303 |
| Hydantoinase | 2.749 | 7.143 | 6.32 | 4.976 | 4.343 | 6.266 | 4.906 | 5.298 | 5.714 | 3.411 | 1.238 | 0.58 | 2.604 | 16.986 | 0.994 | 6.97 | 3.037 | 5.203 | 4.269 | 1.273 | 0.968 | 1.661 | 296.34 | 0.942 | 1.109 |
| T2SSP | 1.309 | 2.207 | 1.022 | 11.251 | 26.172 | 0.186 | 2.207 | 0.598 | 0.371 | 1.347 | 1.273 | 1.109 | 0.274 | 0.261 | 0.635 | 0.288 | 0.616 | 1.05 | 1.139 | 0.844 | 1.079 | 0.545 | 2.303 | 158.136 | 0.821 |
| 2H_CoA_red | 1.022 | 1.309 | 0.511 | 1.772 | 1.715 | 1.171 | 2.749 | 1.966 | 20.03 | 3.963 | 1.079 | 3.194 | 1.772 | 1.427 | 1.109 | 0.673 | 0.916 | 0.462 | 1.609 | 1.109 | 1.833 | 2.501 | 0.301 | 1.204 | 201.038 |

Table S4. Strains and Plasmids Used in This Study

| Strain | Genotype | Source |
|---|---|---|
| yPS003 | W303 (ura3 ade2-1 his3-11,15 leu2-3,112 trp1-1 Mata BUD4 GLK1-sfGFP::spHIS5) | *This Study* |
| yPS031 | W303 (ura3 ade2-1 mCherry::HIS leu2-3,112 trp1-1 Mata BUD4) | *This Study* |
| yPS033 | W303 (ura3 ade2-1 Citrine::HIS leu2-3,112 trp1-1 Mata BUD4) | *This Study* |
| yPS041 | W303 (ura3 ade2-1 Citrine::HIS leu2-3,112 trp1-1 Mata BUD4 glk1Δ::HPH) | *This Study* |
| yPS106 | W303 (ura3 ade2-1 his3-11,15 leu2-3,112 trp1-1 Mata BUD4 Glk1(F3S)) | *This Study* |
| yPS108 | W303 (ura3 ade2-1 his3-11,15 leu2-3,112 trp1-1 Mata BUD4 GLK1(F3S,K182A)) | *This Study* |
| yPS109 | W303 (ura3 ade2-1 his3-11,15 leu2-3,112 trp1-1 Mata BUD4 Glk1(K182A)) | *This Study* |
| yPS110 | W303 (ura3 ade2-1 his3-11,15 leu2-3,112 trp1-1 Mata BUD4 Glk1(F3S)-GFP::URA) | *This Study* |
| yPS111 | W303 (ura3 ade2-1 his3-11,15 leu2-3,112 trp1-1 Mata BUD4 GLK1(F3S,K182A)-GFP::URA) | *This Study* |
| yPS112 | W303 (ura3 ade2-1 his3-11,15 leu2-3,112 trp1-1 Mata BUD4 Glk1(K182A)-GFP::URA) | *This Study* |
| yPS113 | W303 (ura3 ade2-1 Citrine::HIS leu2-3,112 trp1-1 Mata BUD4 Glk1(F3S)) | *This Study* |
| yPS116 | W303 (ura3 ade2-1 Citrine::HIS leu2-3,112 trp1-1 Mata BUD4 Glk1(K182A)) | *This Study* |
| yPS117 | W303 (ura3 ade2-1 Citrine::HIS leu2-3,112 trp1-1 Mata BUD4 Glk1(F3S,K182A)) | *This Study* |

| Plasmid | Description | Source |
|---|---|---|
| pSUMO-Glk1 | his6-SUMO tagged *S. cerevisiae* Glucokinase in T7 expression vector (AMP) | *This Study* |
| pSUMO-caglGlk1 | his6-SUMO tagged *C. glabrata* Glucokinase in T7 expression vector (AMP) | *This Study* |
| pSUMO-caalGlk1 | his6-SUMO tagged *C. albicans* Glucokinase in T7 expression vector (AMP) | *This Study* |
| pSUMO-scpoHxk1 | his6-SUMO tagged *S. pombe* Glucokinase in T7 expression vector (AMP) | *This Study* |
| pSUMO-nacaGlk1 | his6-SUMO tagged *N. castellii* Glucokinase in T7 expression vector (AMP) | *This Study* |
| pSUMO-laklGlk1 | his6-SUMO tagged *L. kluyveri* Glucokinase in T7 expression vector (AMP) | *This Study* |
| pSUMO-vapoGlk1a | his6-SUMO tagged *V. polysporus* Glucokinase-1 in T7 expression vector (AMP) | *This Study* |
| pSUMO-vapoGlk1b | his6-SUMO tagged *V. polysporus* Glucokinase-2 in T7 expression vector (AMP) | *This Study* |
| pSUMO-kllaGlk1 | his6-SUMO tagged *K. lactis* Glucokinase in T7 expression vector (AMP) | *This Study* |
| pSUMO-ergoGlk1 | his6-SUMO tagged *E. gossypii* Glucokinase in T7 expression vector (AMP) | *This Study* |
| pSUMO-Hxk1 | his6-SUMO tagged *S. cerevisiae* Hexokinase-1 in T7 expression vector (AMP) | *This Study* |
| pSUMO-Hxk2 | his6-SUMO tagged *S. cerevisiae* Hexokinase-2 in T7 expression vector (AMP) | *This Study* |
| pSUMO-scpoHxk2 | his6-SUMO tagged *S. pombe* Hexokinase in T7 expression vector (AMP) | *This Study* |

### Movie captions

**Movie S1: Glk1-GFP Polymerization is Induced by Glucose.** Glk1-GFP cells grown to stationary phase were loaded into a flow cell. At the start of the movie, CSM-Glucose medium is flowed over the cells, inducing Glk1 polymerization. The movie is a maximum intensity projection of a confocal z-stack. Images were taken at 1-minute intervals.

**Movie S2: Glk1-GFP Depolymerizes When Glucose is Removed.** Glk1-GFP cells grown to saturation were loaded into a flow cell and washed into glucose containing medium. At the start of the movie, glucose is washed out by washing the cells into CSM medium with no carbon source. The movie is a single confocal slice with continuous capture. Images were taken at 100ms intervals.
