## Supplementary figures and images for "Independent evolution of polymerization in the Actin ATPase clan regulates hexokinase activity"

### Figure S11

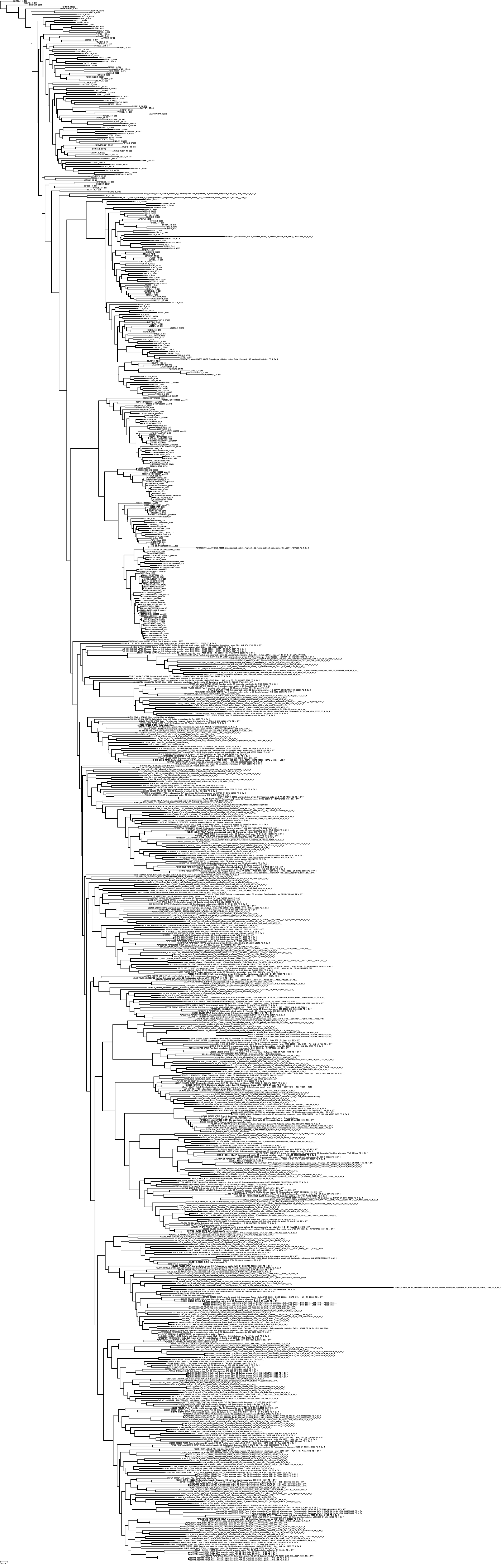

### Table S3

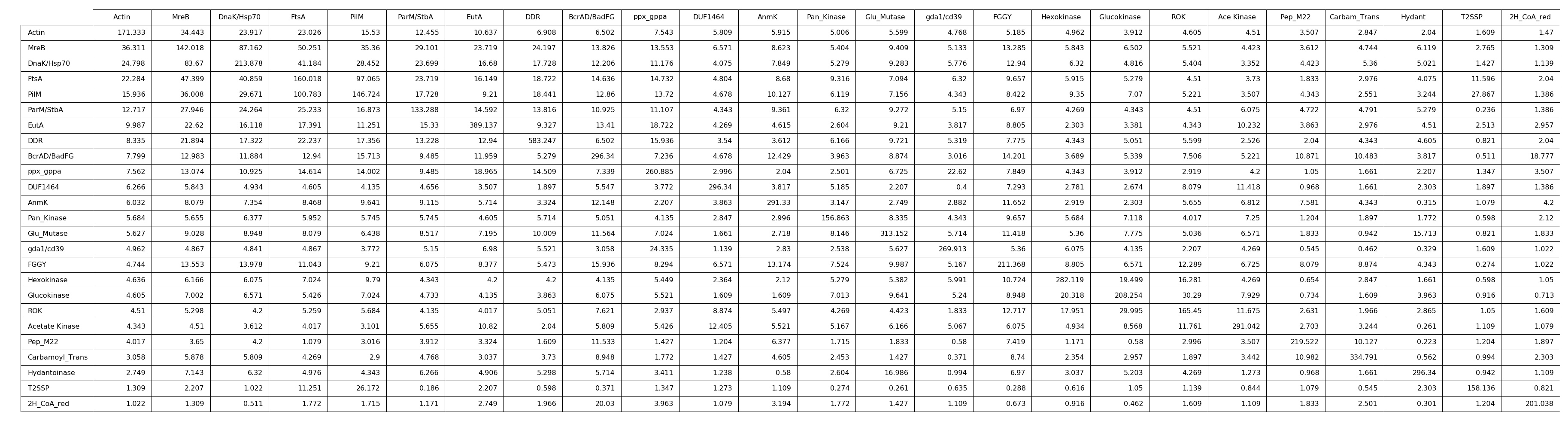
